# Ecological fitness cost associated with the AHAS Trp574Leu mutation in feral *Raphanus sativus*

**DOI:** 10.1101/2020.08.27.270777

**Authors:** Roman B. Vercellino, Fernando Hernández, Claudio E. Pandolfo, Miguel Cantamutto, Alejandro Presotto

## Abstract

Gene mutations endowing herbicide resistance may have negative pleiotropic effects on plant fitness. Quantifying these effects is critical for predicting the evolution of herbicide resistance and developing management strategies for herbicide-resistant weeds. This study reports the effects of the acetohydroxyacid synthase (AHAS) Trp574Leu mutation throughout the life cycle of the weed feral radish (*Raphanus sativus* L.). Resistant and susceptible biotypes responded differently to light and water treatments in relation to germination and emergence. Under light exposure, the resistant biotype showed higher germination and emergence, but no differences were found in seed dormancy, germination in darkness and emergence from buried seeds or pods. The resistant biotype showed delayed and reduced seedling emergence relative to the susceptible biotype under rainfed conditions, but these differences between the biotypes were not detected in irrigated soil. The biotypes showed similar relative growth rates and vegetative biomass. However, under wheat interference, resistant plants had 36–46% less total above-ground biomass, 26–47% less seeds per plant, and 36–53% less plant yield than susceptible ones, and these differences were more evident at higher plant density. This study provides a better understanding of the ecological fitness cost associated with the AHAS Trp574Leu mutation in feral *R. sativus*. The fitness costs could reduce the frequency of the resistant allele in areas untreated with AHAS inhibiting herbicides.

## 1 INTRODUCTION

Acetohydroxyacid synthase (AHAS), also known as acetolactate synthase (ALS), is the first enzyme in the biosynthesis of the branched chain amino acids valine, leucine and isoleucine, and the common target-site of AHAS-inhibiting herbicides (hereafter AHAS herbicides) (Duggleby *et al*., 2008). Repeated use of AHAS herbicides in agricultural fields has resulted in the evolution of AHAS-resistant biotypes in at least 165 weed species worldwide (Heap, 2020). Target-site resistance (TSR) is the most common mechanism resulting in resistant to AHAS herbicides (Yu and Powles, 2014; Heap, 2020). In the AHAS gene, at least 29 resistance-endowing gene mutations in eight conserved amino acids were identified in field-evolved AHAS-resistant weed biotypes (Heap, 2020). Some of them may alter normal enzymatic activity, resulting in a whole plant fitness cost (Vila-Aiub, 2019).

Mutations conferring herbicide resistance, which enable adaptation to a new environment, may have negative pleiotropic effects, also called fitness costs (Vila-Aiub *et al*., 2009). These fitness costs could influence the persistence and evolution of the adaptive alleles in the absence of the selective agent (Baucom, 2019; Vila-Aiub, 2019). It has been reported that fitness costs associated with TSR to AHAS herbicides can be manifested in different life-history traits (i.e. dormancy, germination, early growth and reproductive outputs) (Vila-Aiub *et al*., 2009; Darmency *et al*., 2017) and differentially expressed depending on the specific mutation (Yu *et al*., 2010), weed species (Tardif, *et al*., 2006; Li *et al*., 2013) and environmental conditions (Ashigh and Tardif, 2009; Panozzo *et al*., 2017). There is also a general idea that fitness costs may be more evident under stressful environmental conditions (Vila-Aiub *et al*., 2009; Cousens and Fournier-Level, 2018), but this prediction has not always been identified (Ashigh and Tardif, 2009; Panozzo *et al*., 2017). A case-by-case analysis is thus required to study the expression of the specific gene AHAS-resistant mutation in fitness costs. Quantifying fitness costs associated with herbicide resistant alleles throughout the entire life cycle is necessary to predict the dynamic of the evolution of herbicide resistance and to develop management strategies for herbicide-resistant weeds (Cousens and Fournier-Level, 2018; Baucom, 2019).

*Raphanus sativus* L. (radish) is an ancient root crop from the Brassicaceae family. Spontaneous populations of radish found in Europe, North America, South America and Japan have been classified as de-domesticated (feral) forms derived from the cultivated biotype (Snow and Campbell, 2005). Radish and its feral forms are self-incompatible and insect-pollinated with an annual or biennial life cycle (Snow and Campbell, 2005). Feral radish is a widespread problematic winter weed in temperate zones of the Americas that has developed resistance to AHAS herbicides in Brazil, Chile and Argentina (Pandolfo *et al*., 2016; Heap, 2020). In Argentina, feral radish is one of the most noxious weeds in the south of Buenos Aires province, the main winter cereal producing area (Scursoni *et al*., 2014), and the Trp574Leu mutation has been identified in several AHAS-resistant populations (Pandolfo *et al*., 2016; Vercellino *et al*., 2018).

In a previous study, AHAS-resistant feral radish carrying the Trp574Leu mutation showed negative pleiotropic effects on AHAS activity and reproductive traits under intraspecific interference (Vercellino *et al*., 2018). However, the lack of information of fitness costs associated with the Trp574Leu mutation in different life-history traits of feral radish and under crop interference limits the development and implementation of evolution-knowledge-based resistance management strategies (Cousens and Fournier-Level, 2018). The comparison of resistant and susceptible individuals with similar genetic backgrounds is crucial for the quantification of fitness costs associated with herbicide resistant alleles (Vila-Aiub *et al*., 2011). Here, we compare AHAS-resistant Trp574Leu homozygous individuals with susceptible individuals from a single population to minimize the effect of genetic background (Vila-Aiub *et al*., 2011; Keshtkar *et al*., 2019). The aims of this study were to investigate the effect of the Trp574Leu mutation on (1) seed dormancy, germination and emergence under contrasting conditions of temperature, light, water and burial depth, and on (2) relative growth rate, and vegetative and reproductive traits, under intra- and interspecific interference. This study will provide a better understanding of the fitness costs of the AHAS Trp574Leu mutation in feral *R. sativus*.

## 2 MATERIALS AND METHODS

### 2.1 Plant material

A well-known feral *R. sativus* population (RSBA3) from the south of Buenos Aires province (Argentina) was used in the study (Pandolfo *et al*., 2013, 2016; Vercellino *et al*., 2018). Recently evolved AHAS–resistant plants carrying the Trp574Leu mutation were collected in a field planted with IMI–resistant oilseed rape after imazethapyr application by the farmer (Pandolfo *et al*., 2013, 2016). From these plants, a purified homozygous sub-population with the Trp574Leu mutation (hereafter R biotype) was generated by selecting at least 15 homozygous resistant plants using a specific Trp574Leu mutation CAPS marker, following the procedure described in Pandolfo *et al*. (2016). Their tested AHAS–susceptible counterparts adjacent to the crop, at less than 50 m away from resistant plants, were used to minimize differences in genetic background (hereafter S biotype) (Vercellino *et al*., 2018). Feral radish is a self-incompatible and insect-pollinated species; so therefore, we can assume similar genetic backgrounds in R and S (Vila-Aiub *et al*., 2011; Keshtkar *et al*., 2019).

To increase the amount of seed and to minimize the maternal environmental effects, at least 15 plants of each biotype (R and S) were used to produce a new generation. To prevent cross-pollination, each biotype was isolated in an individual tent built with a pollen-proof mesh. Open pollination was allowed within each biotype (each tent) by adding hives of honeybees (*Apis mellifera*). At the end of the growing season, mature pods were collected and seeds of a portion of pods were extracted by hand or using a mortar. Seeds and intact pods were stored at room temperature and less than 10% moisture until use. Two-hundred progeny individuals were randomly selected from within each sub-population (R and S), and AHAS-resistance and – susceptibility was verified by applying metsulfuron-methyl at double the recommended rate (X=6 g a.i. ha^−1^).

### 2.2 Effects of dry storage, temperature and light on seed germination

To determine the effect of the Trp574Leu mutation on seed dormancy and germination, isolated seeds and seeds inside intact pods of R and S biotypes stored under laboratory conditions for 0 (freshly harvested), 2, 6 and 12 months were tested for germination in 9-cm-diameter Petri dishes incubated at 14/4, 21/9 and 28/14°C in light/darkness (12/12 h) or in constant darkness for 15 days, according to Vercellino *et al*. (2019). The 14/4, 21/9 and 28/14°C regimes represent the average winter, spring–autumn and summer temperatures in the south of Buenos Aires province. At the end of the experiment, the pods were cut open, and the germinated and non-germinated seeds were counted. A seed was considered germinated when the radicle had emerged. The viability of non-germinated seeds was tested using a 1 % (w/v) tetrazolium chloride solution. As the tetrazolium test on the non-germinated seeds showed >99% viability, all non-germinated seeds were considered as alive. The experiment was conducted as a completely randomized design with four replicates.

### 2.3 Effect of burial depth on seedlings emergence

To determine the effect of the Trp574Leu mutation on seeding emergence at different burial depths, twenty-five isolated seeds and five intact pods (29.8 ± 1.2 seeds per replicate) of R and S biotypes (dry-stored for 2 months) were sown at 0, 1, 8, 12 and 16 cm-depths in 10 L (18 cm diameter × 24 cm height) plastic pots containing a sandy-loam soil with 1.1% organic matter and pH 7.7 (the typical soil in the southwest of Buenos Aires province). Six pots per treatment [two seed treatments (isolated seeds and seed inside intact pods) × two biotypes × five soil-depth] were established in a greenhouse (20 ± 5 °C, natural light), and watered regularly. Seedling emergence was evaluated, and the seedlings were removed periodically (3–4 days intervals) for 85 days. For each experiment, the proportion of emerged seedlings was calculated as the ratio between the sown seeds and the emerged seedlings at the end of the experiment. To determine the percentage of emergence in the seeds inside pod treatments, six replicates of ten pods of each biotype were threshed and their seeds were counted. The experiment was conducted as a completely randomized design with six replicates.

### 2.4 Seedling emergence pattern

An outdoor experiment (38°41′38′′S, 62°14′53′′W) was conducted in the Agronomy Department’s experimental field at the Universidad Nacional del Sur, Bahía Blanca, Argentina, to evaluate the seedling emergence pattern of the R and S biotypes. In mid-summer (6 Feb 2018), 50 isolated seeds and 10 intact pods of R and S biotypes were sown at a 0.5 cm-depth in 10 L plastic pots containing the same soil that the previous experiment, under two soil irrigation. Six pots per treatment [two seed treatments (isolated seeds and seed inside intact pods) × two biotypes × two irrigation (irrigated and non-irrigated)] were established. In the irrigated treatment, the soil was watered to field capacity every 3 – 4 days throughout the experiment, while in the non-irrigated treatment the soil only received water from rainfall. Seedling emergence was monitored weekly for one year. Daily minimum and maximum air temperature and rainfall data were obtained from the National Research Council of Argentina (CONICET), Bahía Blanca (38°39′60′′S, 62°13′58′′W) (Fig. S1). For each experiment, the proportion of emerged seedlings was calculated as above. The experiment was conducted as a completely randomized design with six replicates.

### 2.5 Vegetative growth

In a greenhouse in Bahía Blanca, Argentina, three experiments were performed to evaluate the relative growth rate (RGR) and competitive response of R and S biotypes during early vegetative growth. Both eco-physiological parameters are widely used to evaluate the mexpression of herbicide resistance fitness costs (Tardif *et al*., 2006; Li *et al*., 2013; Vila-Aiub *et al*., 2015). Growth experiments were conducted with isolated plants (physiological fitness cost) and plants growing under two interference environments, intraspecific interference and interspecific interference with wheat (*Triticum aestivum* L.) (ecological fitness cost). In all three experiments, the plants were grown in pots (15 cm diameter × 12 cm height) containing the same soil as above, in a greenhouse (20 ± 5 °C, natural light), watered regularly and fertilized with 6 L ha^−1^ of liquid fertilizer (Xilonen^®^, grade 10-13-10) and 30 kg ha^−1^ urea (grade 46-0-0) until they reached the four-leaf stage. The pots were rearranged regularly to randomize any environmental differences within the greenhouse and distributed spatially to avoid competition for light.

#### 2.5.1 Growth of plants without competition

Seeds of the R and S biotypes were germinated in Petri dishes at 14/28 °C in darkness for 48 h (Vercellino *et al*., 2019). At least 50 germinated seedlings of uniform-size (< 1 cm) of R and S were transplanted into individual pots (one plant per pot). The above-ground biomass of 25 individual plants of each biotype were harvested at two and four-leaf stage. The biomass was oven dried for a week at 60 °C and then weighed for dry biomass estimation. Pots were arranged in a completely randomized design with 25 replicates.

#### 2.5.2 Growth of plants under intraspecific competition

A replacement-series design was used to evaluate which of R or S biotypes was the most competitive. Uniform-size germinated seedlings (< 1 cm) of R and S were transplanted equidistantly at four seedlings per pot. R and S seedlings were grown alone or in mixtures with ratios of (R:S) 4:0 (100:0 %), 3:1 (75:25 %), 2:2 (50:50 %), 1:3 (25:75 %) and 0:4 (0:100 %). The above-ground dry biomass from R and S plants was determined at the four-leaf stage, as above. The pots were arranged in a randomized block design with six replicates.

#### 2.5.3 Growth of plants under interspecific competition

A target-neighbourhood design was used to evaluate the competitive responses of R and S biotypes in competition with wheat. One feral radish plant per pot was subjected to competition from increasing densities of neighboring wheat plants (0, 57, 113, 226, 340 and 453 plants m^−2^, which resulted in 0, 1, 2, 4, 6, and 8 plants pot^−1^). Feral radish seedlings (<1 cm) were transplanted into pots when the wheat plants were at the two-leaf stage (Li *et al*., 2013; Vila-Aiub *et al*., 2015). The above-ground dry biomass of R and S biotypes and wheat plants from each experimental unit (pot) was determined when feral radish control plants (0 wheat plants m^−2^) were at the four-leaf stage, as above. The experiments were arranged in a completely randomized design with six replicates.

### 2.6 Reproductive traits under field condition

Field experiments were conducted to evaluate the total above-ground dry biomass and reproductive traits of R and S biotypes grown under interference with wheat. As feral radish is considered a facultative species with two different cohorts (Snow and Campbell, 2005), the experiments were performed on two representative extreme planting dates. The facultative wheat was planted at the end of autumn [mid-May 2018 (winter wheat), cultivar ACA 360 at 200 plants m^−2^] and at the end of winter [early August 2018 (spring wheat), cultivar KLEIN PROTEO at 350 plants m^−2^]. Each experimental unit was composed of nine rows 1.5 m in length and spaced at 0.2 m apart. Seedlings of the R and S biotypes were established in plastic trays containing potting mix (Grow Mix Terrafertil^®^), grown in a greenhouse (20 ± 5 °C, natural light) and watered daily. At the two-three wheat leaf stage, feral radish seedlings (two to three-leaf stage) of each biotype (R and S) were transplanted into the experimental field between the wheat plants according to the planned weed density. The feral radish densities studied were 0 (control), 4, and 12 plants m^−2^. The trials were drip irrigated complementarily and fertilized with 90 kg ha^−1^ diammonium phosphate at planting and 200 kg ha^−1^ urea at the five–leaf stage. The non-target weeds were regularly removed by hand. The experiment was arranged in a split-plot design with the planting date as the main plot, and the biotype and feral radish densities as sub-plots. Within each planting date, experimental units were arranged in a randomized block design with four replicates.

At the end of the growing season, plant height and branch number were measured in four neighbouring feral radish plants located in the centre of each experimental unit. Then, the above-ground biomass from the feral radish plants were hand-harvested, dried at 60 °C to constant weight and weighed to determine the total dry biomass per plant. After that, pods per plant, seeds per pod, seed weight, seeds per plant and plant yield were estimated following the procedure described in Vercellino *et al*. (2018). In each experimental unit, the wheat yield was estimated by hand-harvesting and hand-threshing three replicates of 0.5 m long in the middle of the three central rows. The wheat yield was standardized to 13.5 % moisture content. Data from feral radish and wheat of each experimental unit were averaged for the statistical analysis.

### 2.7 Statistical analysis

Germination and emergence data were analyzed using generalized linear models (GLM) based on maximum likelihood estimation (ML) with PROC GLIMMIX in SAS software (SAS University Edition, SAS Institute Inc., Cary, NC, USA). For experiment ‘*2*.*2*’, a binomial linear model was fitted with a logit link to germination as the response variable. The models included five fixed effects, i.e. light, storage time, temperature, seed treatment (isolated seeds and seeds inside pods) and biotype, and all the interactions.

Two beta models were adjusted due to the non-normal natural distribution of the emergence proportion data. For the experiment ‘*2*.*3*’, the model included three fixed effects, i.e. seed treatment, burial depth and biotype, and all the interactions. For experiment ‘*2*.*4*’, the emergence was evaluated at five different times after the experiment began (1, 3, 6, 9 and 12 months) and the model included three fixed effects, i.e. irrigation (irrigated and non-irrigated), seed treatment and biotype, and all the interactions.

For experiment ‘*2*.*5*.*1*’, the RGR was estimated using the unbiased formula proposed by Hoffmann and Poorter (2002). The variance (*V*) of RGR was estimated with Venus and Causton (1979). The RGR was compared between R and S biotypes using the Student’s t-test with PROC TTEST in SAS. In addition, *ANOVA* was performed to compare the above-ground dry biomass of the R and S biotypes using PROC GLM. The model included harvest time, biotype, and their interaction. Data were log-transformed to improve the normality and homogeneity of variances before analysis.

For experiment ‘*2*.*5*.*2*’, the relative crowding coefficient (RCC) for above-ground dry biomass was calculated according to the formula proposed by Novak *et al*., (1993) following the procedure described by Tardif *et al*., (2006). An RCC value of 1 indicates that both biotypes have equal competitiveness. RCC < 1 shows superior competitiveness of S over R biotype and RCC > 1 indicates the opposite. The RCC values were compared to 1 using the Student’s t-test with PROC TTEST. In addition, *ANOVA* was performed to compare the above-ground dry biomass in the 4:0 (S) and the 0:4 (R) treatments with PROC GLM.

For experiment ‘*2*.*5*.*3*’, data for the above-ground dry biomass of feral radish grown in competition with wheat was expressed as a percentage of that trait in the absence of competition, and it was analyzed following the procedure described in Li *et al*. (2013) and Vila-Aiub *et al*. (2015). The response of R and S biotypes to competition by neighbouring wheat was analyzed using a hyperbolic non-linear model, which described the response of the feral radish plants to increasing biomass of neighbour wheat plants. Slope of the non-linear regression model was compared between the R and S biotypes using the Student’s t-test with PROC TTEST. The hyperbolic non-linear model was fitted after log-transforming data to improve the assumptions of the regression analysis.

In the experiment ‘*2*.*6*’, *ANOVA* with PROC GLM was performed to investigate differences in the total above-ground dry biomass per plant and per area (m^−2^), plant height, branch number, pods per plant, seeds per pod, seeds per plant, seed weight, plant yield and partitioning of the dry biomass (stems and branches, pods and seeds) between the S and R biotypes. The model included planting date, weed density (4 and 12 plants m^−2^), biotype, all their interactions and the block within planting date. In addition, *ANOVA* was performed to evaluate the effect of feral radish interference on wheat yield. The model included planting date, weed density (0, 4 and 12 plants m^−2^), biotype, all their interactions and the block within planting date. Means were compared using Fisher’s LSD test (*P* ≤ 0.05).

## 3 RESULTS

### 3.1 Effects of dry storage, temperature and light on germination

Analysis of the germination at varying times after harvest revealed significant effects of light (*F*_*1;288*_ = 102.68; *P* <0.0001), seed treatment (*F*_*1;288*_ = 77.57; *P* <0.0001), temperature (*F*_*2;288*_ = 31.05; *P* <0.0001), biotype (*F*_*1;288*_ = 6.70; *P* = 0.0101) and the light by seed treatment interaction (*F*_*1;288*_ = 23.97; *P* <0.0001). However, storage time effect, the five-order interaction, the four-order interactions, the three-order interactions and the rest of the two-order interactions were not significant (Table S1). In light, the R biotype showed higher seed germination than the S throughout the experiment, both inside pods and isolated, especially in the 9/21°C and 14/28°C treatments (Fig. 1A and C). These differences between the biotypes were not detected at low temperatures (4/14 °C), nor in seed germination under darkness, either inside the pods or isolated (Fig. 1B and D).

**Figure 1.**
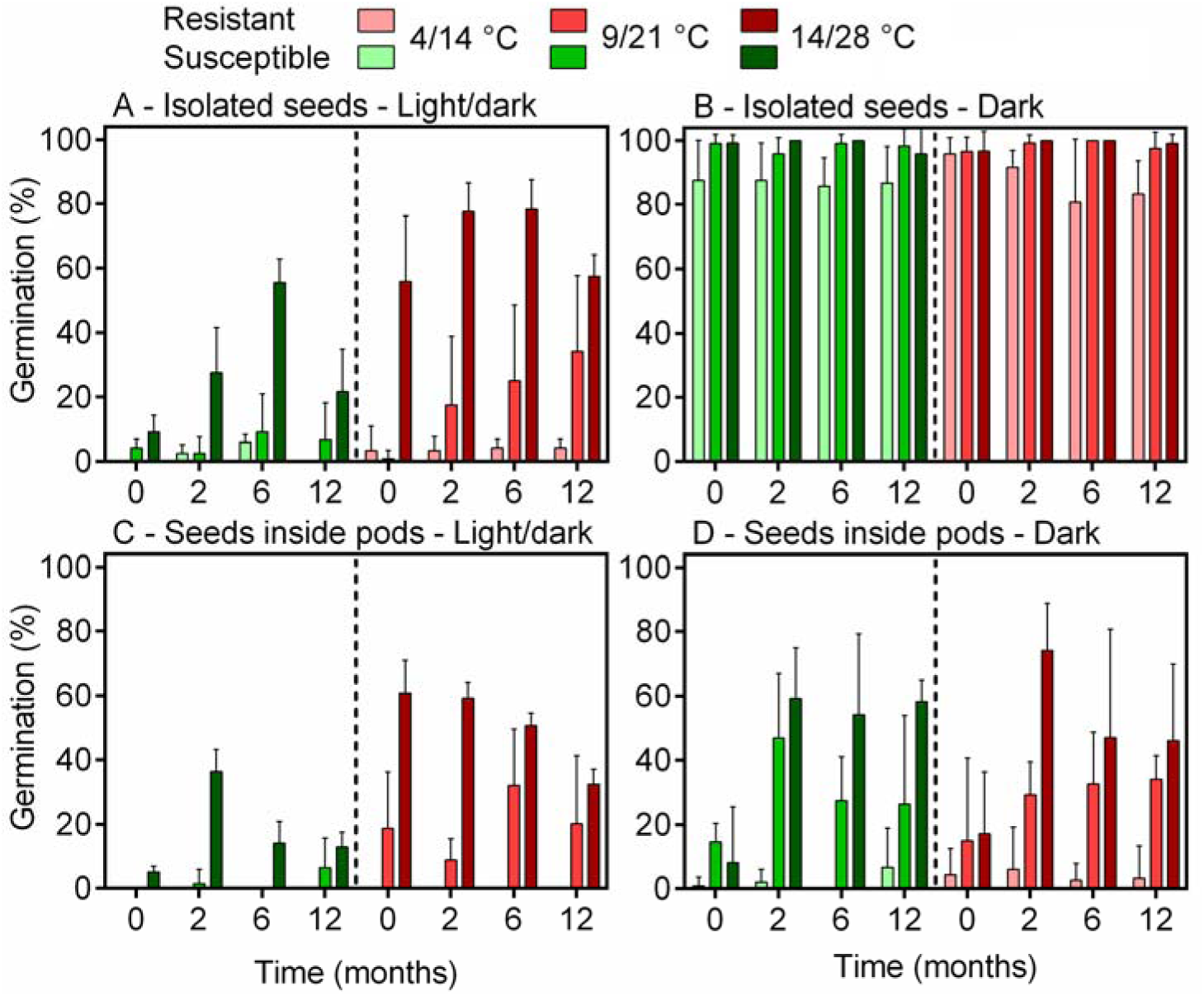
Final germination percentages (means ± 95 % confidence intervals) of isolated seeds under light/dark (A), isolated seeds under constant darkness (B), seeds inside pods under light/dark (C) and seeds inside pods under constant darkness (D) of AHAS herbicide-resistant Trp475Leu (R) and susceptible wild-type (S) feral *Raphanus sativus* biotypes, incubated at three alternating temperature regimes following 0, 2, 6 and 12 months of dry storage under laboratory conditions.

### 3.2 Effect of burial depth on seedlings emergence

The analysis of the total seedling emergence revealed significant effects of the seed treatment × burial depth × biotype three-order interaction (*F*_*5;120*_ = 2.58; *P* = 0.0299). Therefore, we compared the differences in seedling emergence between the R and S biotypes for each seed treatment and burial depth. On the soil surface, isolated seeds of the R biotype showed 79% (25 vs. 14%) higher germination than the S (*F*_*1;10*_ = 8.73; *P* = 0.0144) (Fig. 2A). Inside pods, less than 1% of the seeds germinated on the soil surface in both biotypes (Fig. 2A). As the burial depth increased, seedling emergence from both the isolated seeds and intact pods decreased, except for seeds sown on the soil surface. Seedling emergence from isolated seeds was higher than from the intact pods in both biotypes at all the depths evaluated (Fig. 2), except at 16 cm where less than 1% emergence was detected in both the isolated seeds and the intact pods (data not shown). However, no significant differences were found between the biotypes in cumulative seedling emergence when isolated seeds and intact pods were below the soil surface at up to 16 cm deep (Fig. 2B, C and D).

**Figure 2.**
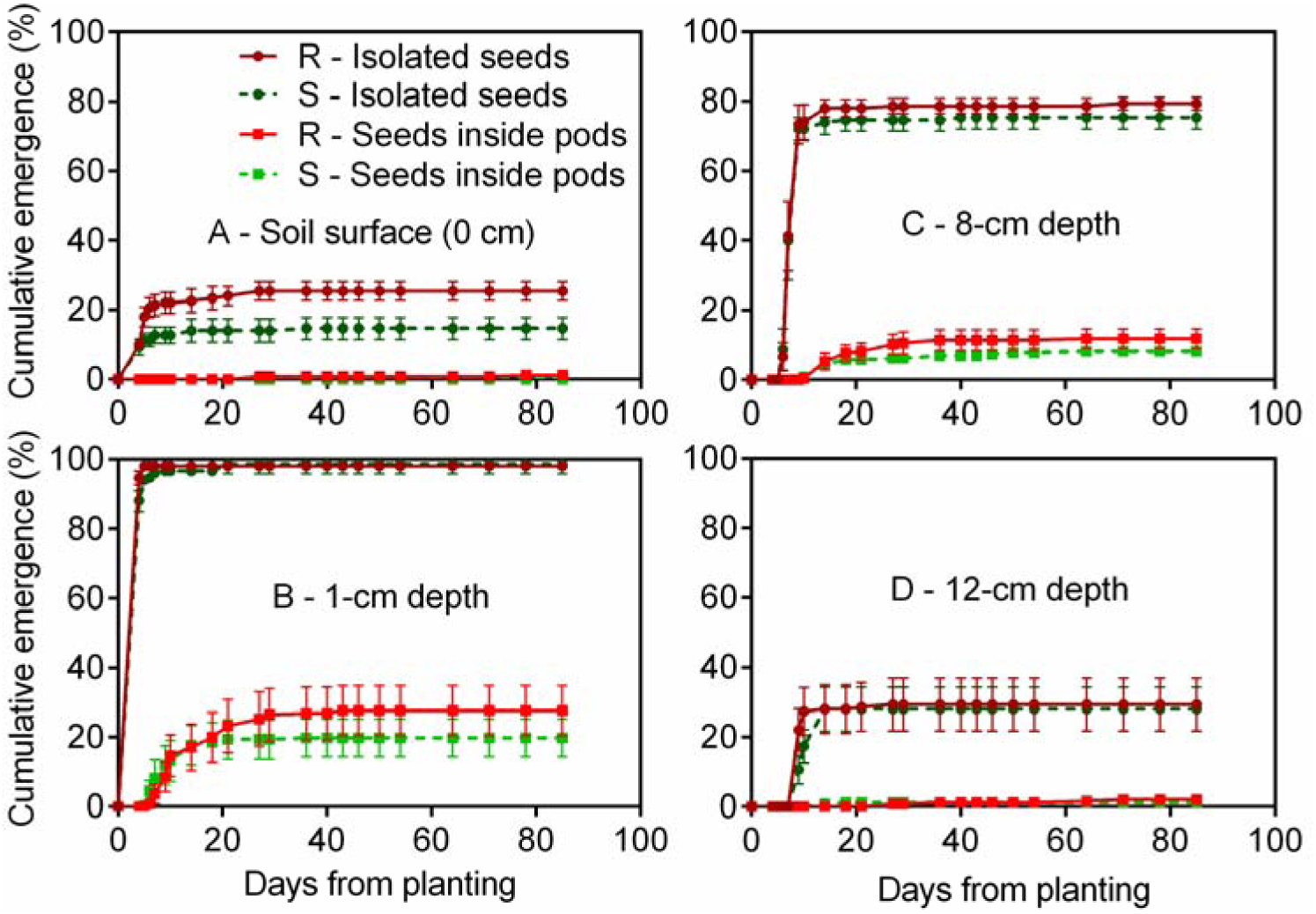
Dynamics of seedling emergence from isolated seeds and seeds inside pods, buried at 0 (A), 1 (B), 8 (C) and 12 (D) cm depth, of AHAS herbicide-resistant Trp475Leu (R) and susceptible wild-type (S) feral *Raphanus sativus* biotypes. Vertical bars represent the standard error of the mean.

### 3.3 Seedling emergence pattern

The mean annual temperature was 15.4°C and mean monthly maximum and minimum temperatures of the coldest (July) and hottest (January) months were 12.0 and 3.0°C (mean 7.1 °C) and 32.6 and 16.2°C (mean 23.3°C), respectively (Fig. S1). Annual precipitation was 592 mm, about 25% (152 mm) in autumn, 11% (65 mm) in winter, 39% (229 mm) in spring and the remaining 25% (89 and 58 mm) in summer (6-Feb-2018 – 20-Mar-2018 and 21-Dec-2018 – 6-Feb-2019, respectively) (Fig. S1).

Analysis of seedling emergence revealed significant effects of the irrigation by seed treatment and irrigation by biotype interactions throughout the study period (Table S2). Therefore, we compared the differences in seedling emergence between the R and S biotypes for each irrigation condition (Table S3). The biotype by seed treatment interaction was not significant for any irrigation condition throughout the experiment (Table S3). The presence of the pericarp strongly reduced seedling emergence in both the irrigation conditions and biotypes throughout the study period (Fig. 3, Table S3).

**Figure 3.**
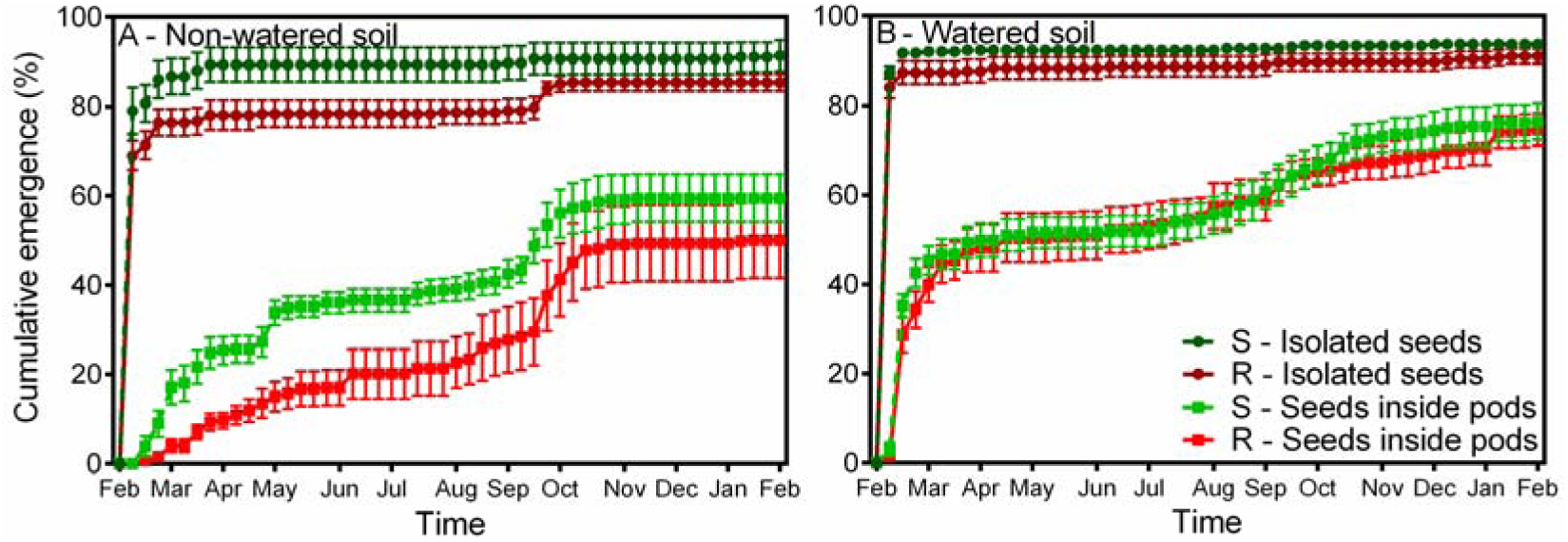
Dynamics of seedling emergence from isolated seeds and seeds inside pods, buried at 1 cm depth in non-irrigated (A) and irrigated (B) soil conditions, of AHAS herbicide-resistant Trp475Leu (R) and susceptible wild-type (S) feral *Raphanus sativus* biotypes. Vertical bars represent the standard error of the mean.

Under rainfed conditions, the R biotype delayed and reduced seedling emergence relative to the S (Fig. 3, Table S3). Seedling emergence of the R biotype was 13% (76 vs. 87%) and 76% (4 vs. 17%) lower than of the S biotype, from isolated seeds and intact pods, respectively, during the first month (Fig. 3A). The R biotype showed a 6% additional emergence from isolated seeds at the end of winter (Fig. 3A). The R and S biotypes showed 16 and 20% emergence during the autumn and 28 and 20% emergence at the end of winter and in early spring, respectively (Fig. 3A). At the end of the experiment, the R biotype showed 7% (91 vs. 85%) and 15% (59 vs. 50%) lower seedling emergence than the S from isolated seeds and intact pods, respectively (Fig. 3A, Table S3).

No differences were detected between the R and S biotypes in irrigated soil (Fig. 3B, Table S3). Under this condition, about 90% of seedlings emerged from isolated seeds and only 40-45% from intact pods during the first month (Fig. 3B). Seedling emergence from intact pods exceeded 50% during autumn (mid-May) and raised an additional 25% between late winter and spring (Fig. 3B). At the end of the experiment, the total seedling emergence was more than 90% in the isolated seeds and 75% in the intact pods, with no differences between the biotypes (Fig. 3B, Table S3).

### 3.4 Vegetative growth under isolation, intra and interspecific competition

No significant differences (*t* = 0.76; *P* = 0.45) were found in the RGR between R (0.1206 ± 0.0047 mg mg^−1^ día^−1^) and S (0.1258 ± 0.0048 mg mg^−1^ día^−1^) biotypes. There was no significant harvest time by biotype interaction (*F*_1;96_ = 0.29; *P* = 0.59). Dry biomass of the R biotype at two (295.02 ± 15.35 mg plant^−1^) and four (1672.55 ± 144.65 mg plant^−1^) -leaf stage did not show any significant differences (*F*_1;96_ = 3.23; *P* = 0.07) compared to the S plants at two (322.37 ± 17.4 mg plant^−1^) and four (1935.36 ± 143.73 mg plant^−1^) -leaf stage.

In the absence of inter-biotype competition, treatments 4:0 and 0:4, no significant differences (*F*_1;11_ = 0.64; *P* = 0.44) were found in the dry biomass between S and R (Fig. 4). However, when R and S biotypes were grown in a mixture, the RCC value was 1.25 (*t* = 2.96; *P* = 0.0253), indicating a slight competitive advantage of S relative to R (Fig. 4).

**Figure 4.**
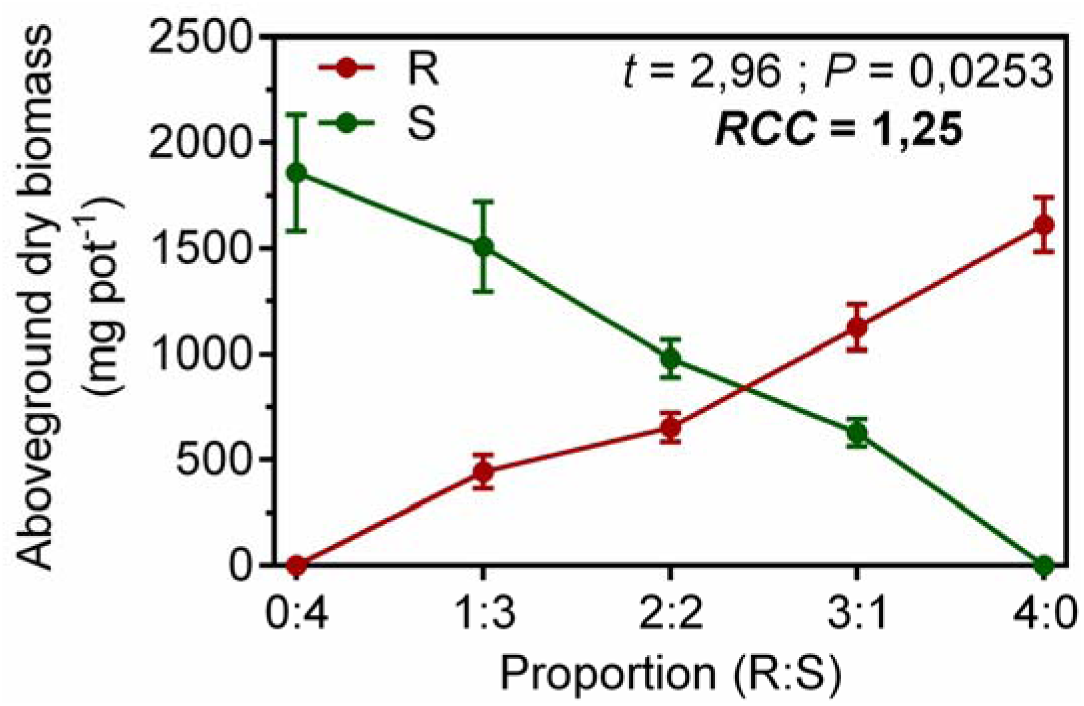
Replacement series diagram for the above-ground vegetative dry biomass of AHAS herbicide-resistant Trp475Leu (R) and susceptible wild-type (S) feral *Raphanus sativus* biotypes 35 days after transplanting. Vertical bars represent the standard error of the mean.

The hyperbolic model adequately explained variations in the feral radish dry biomass to increasing biomass of neighboring wheat plants (*R*^*2*^ = 0.57 – 0.58, *P* <0.0001). This led to a reduction in the feral radish dry biomass with increasing wheat competition (Fig. 5). Competitive responses, evaluated by comparing estimates of regression slopes between the R and S biotypes in the presence of wheat, did not show any significant differences (*P* > 0.95), indicating similar reductions in feral radish dry biomass in the R and S biotypes with increasing wheat biomass (Fig. 5).

**Figure 5.**
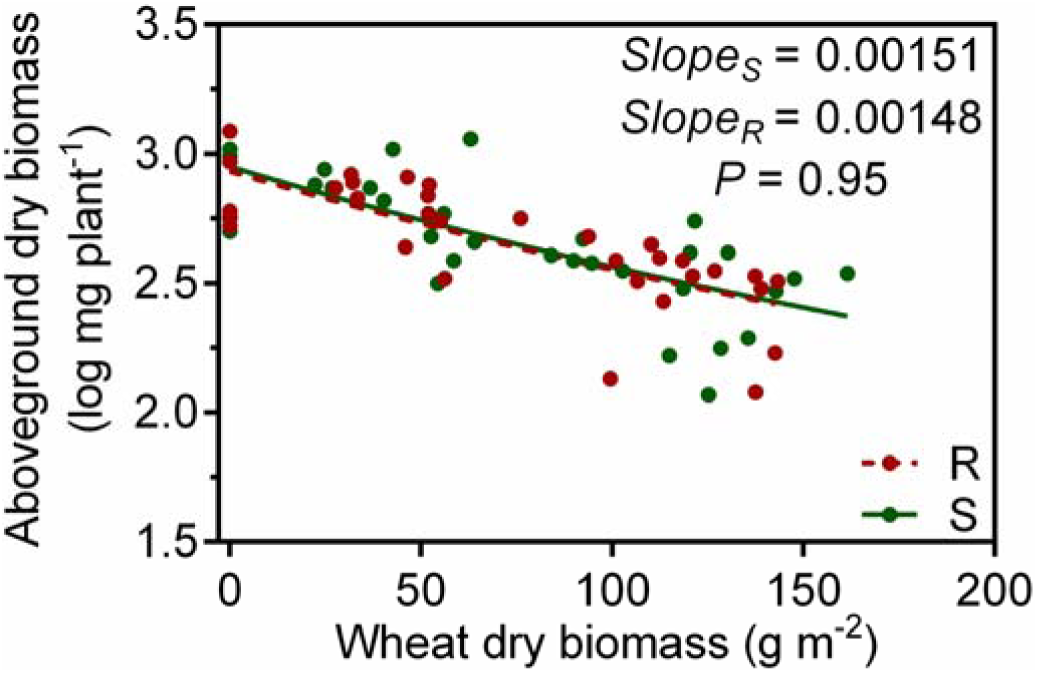
Variations in the above-ground vegetative dry biomass of target AHAS herbicide–resistant Trp475Leu (R) and susceptible wild-type (S) feral *Raphanus sativus* individuals under increasing biomass of wheat neighbour plants. Comparison of regression slopes (*b* parameter) determines differences in competitive responses of the target plants to wheat neighbour plants.

### 3.5 Evaluation of reproductive traits under field conditions

There were no significant interactions between planting date and biotype, planting date and density, biotype and density and the third-order interaction for all evaluated traits, therefore data from the two planting dates were pooled (Table S4, Fig. 6). Significant differences between S and R biotypes were found for the total aboveground biomass (*F*_*1;31*_ = 37.0; *P* <0.0001), pod number (*F*_*1;31*_ = 16.2; *P* = 0.0008), seed number (*F*_*1;31*_ = 27.5; *P* <0.0001), seed weight (*F*_*1;31*_ = 19.0; *P* = 0.0004) and plant yield (*F*_*1;31*_ = 51.05; *P* <0.0001), but no significant differences were found in plant height, branch number and seeds per pod (Table S4). The R biotype had 36–46% less total dry biomass, 20–48% fewer pods per plant, 10–11% less seed weight, 26–47% fewer seeds per plant, and 36–53% lower plant yield than its counterpart S, for both planting dates and weed densities (Table S4, Fig. 6).

**Figure 6.**
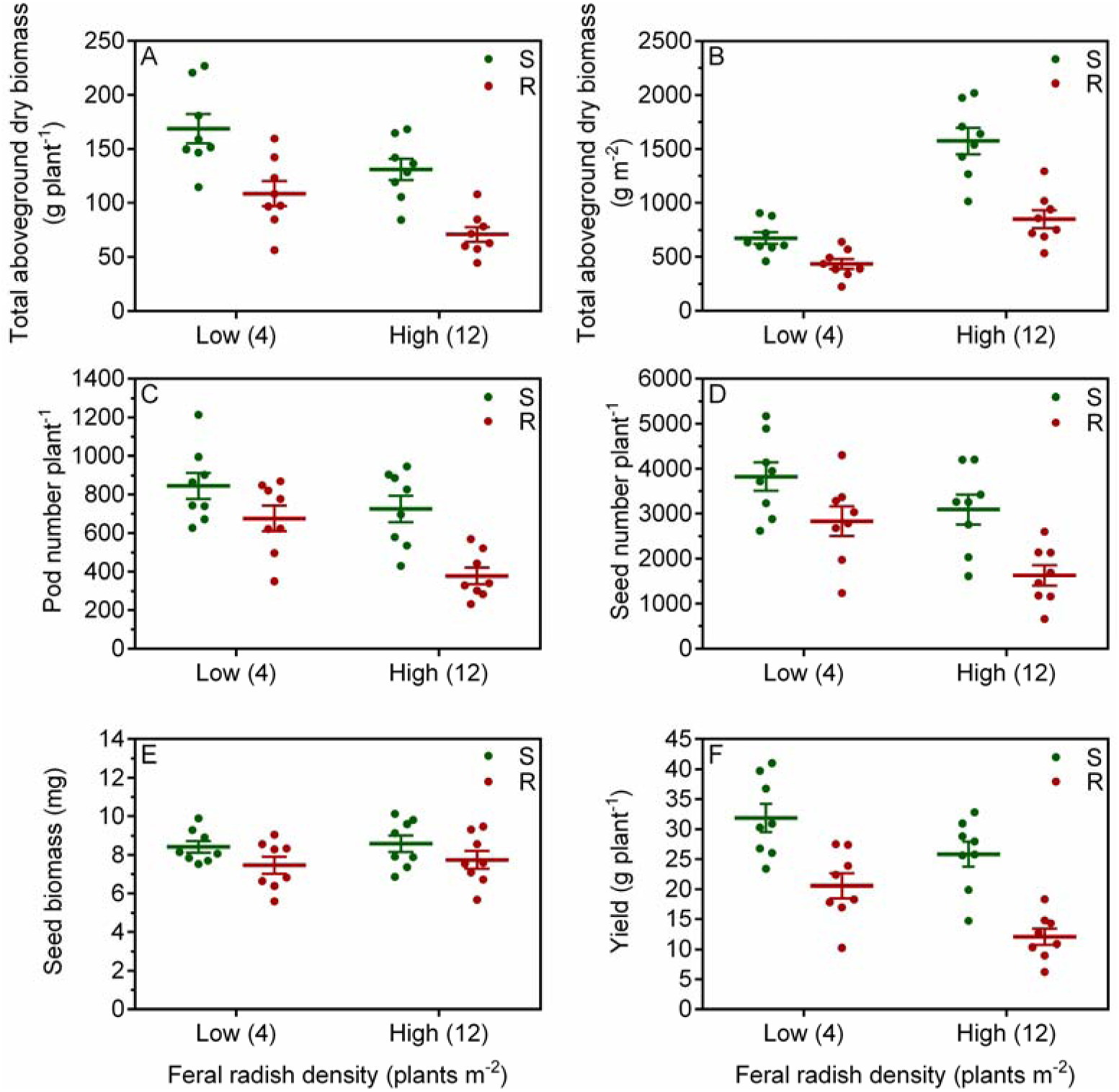
Total above-ground dry biomass per plant (A) and per area (B), pods per plant (C), seeds per plant (D), seed biomass (E) and plant yield (F) of AHAS herbicide-resistant Trp475Leu (R) and susceptible wild-type (S) feral *Raphanus sativus* biotypes. Data are the average of two planting dates. Each point (●;●)is the average of four plants. Vertical bars represent the standard error of the mean.

Analysis of biomass allocation to reproductive traits (i.e. proportion of seed biomass relative to total dry biomass) revealed significant effects of biotype by density, and biotype by planting date interactions (Table S4). Therefore, the data were separated by planting date and weed density (Fig. S2). At 4 weed plants m^−2^, no significant differences were found in the biomass allocation to reproductive structures between the R and S biotypes, neither in winter (*F*_*1; 7*_ = 3.06; *P* = 0.18) or in spring (*F*_*1;7*_ = 0.63; *P* = 0.49). However, at 12 weed plants m^−2^, the R plants showed 19 % (18.8 ± 0.5 vs. 15.2 ± 0.6 %) and 9 % (20.4 ± 0.6 vs. 18.5 ± 0.6 %) lower biomass allocation to reproduction structures than the S plants in both winter (*F*_*1;7*_ = 24.75; *P* = 0.0156) and spring (*F*_*1;7*_ = 14.20; *P* = 0.0327) (Fig. S2).

The wheat yield was significantly reduced by feral radish interference at both weed densities and biotypes (Table S5). No significant differences were found between planting dates, block (inside each planting date) and all the interactions (Table S5). The interference of the R biotype reduced the wheat yield less than the S biotype at both feral radish densities (Fig. S3). The interference of R and S feral radish biotypes reduced the wheat yield by 38 and 49 % at 4 weed plants m^−2^ and 54 and 66 % at 12 weed plants m^−2^, respectively (Fig. S3).

## 4 DISCUSSION

In the present study, we evaluated several life-history traits defining the overall fitness costs associated with the AHAS Trp574Leu mutation in feral radish. In relation to germination and emergence, no differences were found between the R and S biotypes in seed dormancy and germination in darkness, or in seedling emergence from buried seeds/pods, but the R biotype showed higher germination and emergence than the S under light exposure (Fig. 1 and 2). The R biotype showed delayed and reduced seedling emergence relative to the S under rainfed conditions, but these differences were not detected under irrigation (Fig. 3). Finally, both biotypes showed similar relative growth rates and vegetative biomass. However, the R biotype showed lower competitive ability, total above-ground dry biomass and reproductive outputs than the S (Fig. 4 and 6). All of these suggest that the pleiotropic effects imposed by the Trp574Leu mutation can be expressed throughout the life-history of feral radish and in response to several conditions, such as light exposure and water availability, highlighting the importance of considering both the whole life-history and environmental variation in fitness cost studies.

### 4.1 Seed germination and seedling emergence

We previously reported that fresh isolated seeds of feral radish were non-dormant, but that light and the pericarp reduced germination, especially under low temperatures (Vercellino *et al*., 2019), as was observed here (Fig. 1). This is consistent with the high emergence from buried isolated seeds and reduced emergence from seeds inside pods and seeds/pods on soil surface (light exposure) (Fig. 2). When seeds and pods were buried, seedlings emergence decreased with increasing burial depth, and this effect was greater when the seeds were inside pods. In darkness, no differences were found in seed germination between the R and S biotypes, isolated or inside pods, or in the emergence from buried seeds/pods. So, the darkness condition together with the release of the seeds from the pod by breaking the pericarp – e.g. during sowing or tillage up to 12-cm-deep – would promote germination and emergence of feral radish (Chauhan *et al*., 2006; Vercellino *et al*., 2019). Under these conditions, there would be no differences in seedling emergence between the biotypes.

However, the light and irrigation conditions showed differential responses between the R and S biotypes. Light stimulated an increase in seed germination, whether isolated or inside pods, in the R biotype. Accordingly, the R biotype showed higher emergence than the S in isolated seeds placed on the soil surface. The increased emergence may have an adaptive importance in agricultural systems resembling no-till systems, widely used in Argentina (>90%) (Scursoni *et al*., 2019), where weed seeds are often exposed on the soil surface, e.g. after pod scarification by the harvester. The seeds inside pods of both biotypes germinated <2% on the soil surface at least five months after the beginning of the experiment (data not shown), suggesting the development of secondary dormancy, which could be crucial for seed persistence in the soil seedbank (Baskin and Baskin, 2014; Vercellino *et al*., 2019).

Under outdoor conditions, seedling emergence from isolated seeds was concentrated at the end of the summer, soon after seed or pod dispersal and the presence of the pericarp (seeds inside pods) reduced, spread and/or prolonged the seedling emergence period, as was suggested previously (Vercellino *et al*., 2019). Seedling emergence from intact pods was mainly concentrated in autumn and spring in both biotypes, independent of water availability, possibly associated with medium/high temperatures, and with higher precipitations under rainfed conditions. In addition, high humidity and medium/high temperatures could have increased the growth potential of the embryo and/or reduced the mechanical resistance of the pericarp, due to its softening or degradation by water and/or fungal activity (Sperber *et al*., 2017; Steinbrecher and Leubner-Metzger, 2017; Vercellino *et al*., 2019). No differences were found between the biotypes in the seedling emergence pattern under irrigation, whereas the R biotype showed delayed and reduced seedling emergence relative to the S under rainfed conditions. It has been suggested that selection for late germinating seed cohorts is associated with herbicide resistance (reviewed in Darmency *et al*. (2017)), and although the genetic or physiological basis of this trait is unknown, it could be linked to an increase in or development of dormancy (Baskin and Baskin, 2014; Darmency *et al*., 2017). For example, it has been shown an increase in seed dormancy selected along with herbicide resistance in high-intensity cropping systems (Owen *et al*., 2011). The possibility that the effects observed here were caused by environmental selection instead of the pleiotropic effects of the Trp574Leu mutation cannot be ruled out. Additional experiments would be necessary to check our inferences and the biological, ecological and evolutionary significances of these results.

### 4.2 Plant fitness: Vegetative and reproductive traits

In the present study, R and S isolated individuals, or those under intra-/interspecific competition, had similar relative growth rates and vegetative biomass. These results are in agreement with two previous studies showing similar relative growth rates and competitive resource response associated with the Trp574Leu mutation in *Raphanus raphanistrum* (Li *et al*., 2013) and *Lolium rigidum* (Yu *et al*., 2010). Conversely, the R individuals showed a slight competitive disadvantage during intraspecific competition with the S individuals. A similar result was found in AHAS-resistant *Amaranthus powellii* with the Trp574Leu mutation (Tardif *et al*., 2006). However, in *A. powellii* the relative crowding coefficient value was >6, much higher than that observed in our experiment (RCC = 1.25).

Under field conditions, the R individuals grown with wheat interference had a lower above-ground total biomass than the S wild-type with similar resource allocation to reproductive traits (it was only slightly affected at the highest weed density). This resulted in R individuals with a lower number of pods and seeds per plant, lower seed biomass and less yield per plant compared with the S ones. These results are in agreement with our previous study carried out in the absence of crop interference (Vercellino *et al*., 2018). However, fitness costs found here were slightly higher than that found in the absence of crop interference (Vercellino *et al*., 2018). These studies reveal that fitness costs associated with AHAS Trp574Leu mutant *R. sativus* plants are evident under a wide range of environmental conditions. The physiological mechanism responsible for fitness costs associated with the Trp574Leu mutation in *R. sativus* is likely to be associated with the adverse effect of this mutation on AHAS activity relative to the S wild-type (Vercellino *et al*., 2018). Consequently, reduced AHAS activity due to the Trp574Leu mutation could be expressed under resource limiting conditions or in advanced stages of weed life-history. Accordingly, S individuals showed higher wheat yield losses than the R at both weed densities and planting dates.

### 4.3 Eco-evolutionary implications of AHAS herbicide resistance

The ecological and evolutionary implications of an adaptive mutation, e.g. the AHAS Trp574Leu mutation, depends on multiple interactions between operational factors related to herbicide application, genetic factors related to fitness benefits/costs and biological factors focused on weed ecology and life-history traits (Vila-Aiub, 2019). In AHAS herbicide-treated areas, it is expected that the Trp574Leu mutation increases their frequency (Pandolfo *et al*., 2016). However, our studies (Vercellino *et al*., 2018) suggest negative pleiotropic effects on the reproductive traits of feral radish imposed by the Trp574Leu mutation, which would reduce the frequency of the R allele in both ruderal and agrestal environments without AHAS herbicide selection (Baucom, 2019; Vila-Aiub, 2019). In cross pollinated species, such as feral radish (Snow and Campbell, 2005), gene flow between R and S populations could modify the dynamics of R alleles (Maino *et al*., 2019). Gene flow from populations with null/low resistance level could act as a refuge, delaying resistance evolution in populations with a high level of herbicide selection and/or decreasing the frequency of R alleles in herbicide untreated areas. Accordingly, the importance of generating lower selection pressure is highlighted and also maintaining fence lines and fields margin without any herbicide application (Maino *et al*., 2019).

The eco-evolutionary consequences of the differential germination and emergence responses between R and S biotypes are environmentally dependent and complex to predict. It is the ecological environment that will determine the advantage or disadvantage of one biotype over the other during seed germination and seedling emergence (see Section 4.1). On the other hand, a persistent feral radish seedbank could act as a refuge, delaying the evolution of resistance or reducing R alleles. All this knowledge could be incorporated into herbicide resistance evolution models to explore the consequences of our results on the dynamics of resistance to AHAS herbicides and identify possible management strategies to prevent, delay or reverse weed resistance evolution.

These results should not be generalized because there could be an interaction between the Trp574Leu mutation and the genetic background (Paris *et al*., 2008; Vila-Aiub, 2019), although a previous study comparing several R and S populations showed similar results (Vercellino *et al*., 2018). The possibility that the effects observed here were caused by genetic linkage or phenotypic correlations with other traits instead of the pleiotropic effects of the Trp574Leu mutation and only correlated with it, cannot be completely ruled out (Darmency *et al*., 2017; Baucom, 2019). To reinforce these results, future research should focus on monitoring the frequency of the AHAS Trp574Leu mutation for several years under field conditions, in the absence of AHAS herbicide selection, using R and S biotypes.

## ACKNOWLEDGEMENTS

This work was supported by grant ANPCYT-PICT 2012-2854. We thank the National Research Council of Argentina (CONICET) for a fellowship to FH and RBV.

