## Supplementary material for "Ecological fitness cost associated with the AHAS Trp574Leu mutation in feral *Raphanus sativus*": Fig. S2 (from another version of this preprint: version 2, 2020-08-31)

Time (months)

Temperature (°C) or raninfall (mm)

|  | Rainfall | T max | T min |
| --- | --- | --- | --- |
| 2018-02-01 | 0 | 27 | 12.6 |
| 2018-02-02 | 0 | 36.5 | 13.7 |
| 2018-02-03 | 0 | 36.2 | 13.5 |
| 2018-02-04 | 0 | 37.4 | 19 |
| 2018-02-05 | 2 | 37.5 | 21.5 |
| 2018-02-06 | 5 | 38.3 | 21.6 |
| 2018-02-07 | 0 | 38.5 | 22.1 |
| 2018-02-08 | 0 | 33 | 20 |
| 2018-02-09 | 0 | 35.8 | 15.8 |
| 2018-02-10 | 0 | 30 | 18 |
| 2018-02-11 | 0 | 22 | 14.6 |
| 2018-02-12 | 0 | 21.3 | 11 |
| 2018-02-13 | 0 | 28.6 | 6.8 |
| 2018-02-14 | 0 | 32.3 | 16.4 |
| 2018-02-15 | 0 | 34.8 | 17.8 |
| 2018-02-16 | 0 | 36.4 | 17.4 |
| 2018-02-17 | 0.5 | 37.4 | 23.1 |
| 2018-02-18 | 3 | 37.8 | 20 |
| 2018-02-19 | 0 | 35.2 | 22 |
| 2018-02-20 | 0 | 28.4 | 16 |
| 2018-02-21 | 13 | 23.8 | 11.5 |
| 2018-02-22 | 15 | 27.2 | 15.3 |
| 2018-02-23 | 0 | 28.5 | 15.8 |
| 2018-02-24 | 0 | 24 | 16.2 |
| 2018-02-25 | 0 | 30.5 | 12 |
| 2018-02-26 | 0 | 32 | 13.4 |
| 2018-02-27 | 0 | 32 | 13.5 |
| 2018-02-28 | 6 | 24.3 | 18 |
| 2018-03-01 | 0 | 30 | 15.8 |
| 2018-03-02 | 0 | 35.1 | 16.2 |
| 2018-03-03 | 0 | 37.2 | 17 |
| 2018-03-04 | 0 | 36.5 | 23 |
| 2018-03-05 | 0 | 33.6 | 20.6 |
| 2018-03-06 | 0 | 23.8 | 12.2 |
| 2018-03-07 | 0 | 30.6 | 7.8 |
| 2018-03-08 | 0 | 32.8 | 16.2 |
| 2018-03-09 | 0 | 34.8 | 16 |
| 2018-03-10 | 0 | 35.8 | 21.6 |
| 2018-03-11 | 38.2 | 30 | 12.8 |
| 2018-03-12 | 0 | 25 | 7.4 |
| 2018-03-13 | 0 | 29.8 | 10.3 |
| 2018-03-14 | 0 | 24.2 | 9.9 |
| 2018-03-15 | 0 | 27 | 10.3 |
| 2018-03-16 | 2 | 23.8 | 5 |
| 2018-03-17 | 0 | 31.1 | 13.6 |
| 2018-03-18 | 0 | 27.3 | 17.9 |
| 2018-03-19 | 4 | 20.8 | 13.4 |
| 2018-03-20 | 0 | 27.6 | 6.5 |
| 2018-03-21 | 0 | 19 | 9 |
| 2018-03-22 | 0 | 22 | 6.6 |
| 2018-03-23 | 0 | 26.5 | 10.3 |
| 2018-03-24 | 6 | 23.4 | 11.7 |
| 2018-03-25 | 2.1 | 18 | 11 |
| 2018-03-26 | 2 | 19.6 | 5.2 |
| 2018-03-27 | 0 | 23.4 | 8.4 |
| 2018-03-28 | 0 | 27 | 10.9 |
| 2018-03-29 | 0 | 31.4 | 16.9 |
| 2018-03-30 | 0 | 23 | 15 |
| 2018-03-31 | 0 | 29 | 9.8 |
| 2018-04-01 | 0 | 29.9 | 12.8 |
| 2018-04-02 | 0 | 26 | 12.6 |
| 2018-04-03 | 0 | 27.9 | 9.6 |
| 2018-04-04 | 0 | 28.4 | 13.7 |
| 2018-04-05 | 0 | 33.8 | 18.1 |
| 2018-04-06 | 0 | 26.2 | 8.8 |
| 2018-04-07 | 0 | 20.5 | 6.6 |
| 2018-04-08 | 0 | 23.8 | 6.5 |
| 2018-04-09 | 0 | 24.5 | 13.3 |
| 2018-04-10 | 0 | 26.1 | 10.8 |
| 2018-04-11 | 0 | 30.2 | 15.3 |
| 2018-04-12 | 10 | 23.8 | 14.5 |
| 2018-04-13 | 0 | 17.9 | 8.2 |
| 2018-04-14 | 0 | 21 | 1.6 |
| 2018-04-15 | 0 | 24.5 | 4.2 |
| 2018-04-16 | 0 | 27.5 | 12.3 |
| 2018-04-17 | 0 | 29.2 | 17 |
| 2018-04-18 | 0 | 24.1 | 13 |
| 2018-04-19 | 0 | 24.9 | 8.4 |
| 2018-04-20 | 0 | 29.2 | 9.2 |
| 2018-04-21 | 0 | 31.4 | 14.5 |
| 2018-04-22 | 0 | 31.6 | 14.8 |
| 2018-04-23 | 12 | 19.3 | 12.9 |
| 2018-04-24 | 0 | 18.8 | 9.8 |
| 2018-04-25 | 29 | 14 | 8.8 |
| 2018-04-26 | 0.9 | 19.6 | 10.6 |
| 2018-04-27 | 0 | 26.4 | 15 |
| 2018-04-28 | 2 | 24.2 | 9.8 |
| 2018-04-29 | 0 | 16.6 | 6.9 |
| 2018-04-30 | 0 | 19.6 | 9.5 |
| 2018-05-01 | 0 | 16.3 | 6.8 |
| 2018-05-02 | 0 | 17 | 4.8 |
| 2018-05-03 | 7 | 15.8 | 8.8 |
| 2018-05-04 | 7 | 18.8 | 11.4 |
| 2018-05-05 | 0.6 | 14.4 | 11 |
| 2018-05-06 | 0 | 19.5 | 6.4 |
| 2018-05-07 | 0 | 23.2 | 11 |
| 2018-05-08 | 6 | 21.5 | 14.8 |
| 2018-05-09 | 14 | 19.2 | 14.5 |
| 2018-05-10 | 26 | 20.2 | 12.8 |
| 2018-05-11 | 5 | 22 | 14.8 |
| 2018-05-12 | 0 | 20 | 7.4 |
| 2018-05-13 | 0 | 17.2 | 4.6 |
| 2018-05-14 | 0 | 21.5 | 5.2 |
| 2018-05-15 | 1 | 18.6 | 10.9 |
| 2018-05-16 | 0 | 14 | 4 |
| 2018-05-17 | 0 | 13.8 | 2 |
| 2018-05-18 | 4 | 16.1 | 8 |
| 2018-05-19 | 2 | 14.4 | 5 |
| 2018-05-20 | 0 | 13.4 | 3.3 |
| 2018-05-21 | 0 | 14.3 | 6.4 |
| 2018-05-22 | 0 | 15 | 3.3 |
| 2018-05-23 | 0 | 15 | 0.2 |
| 2018-05-24 | 0 | 17 | 2 |
| 2018-05-25 | 0 | 21.4 | 0.3 |
| 2018-05-26 | 0 | 20.6 | 10.2 |
| 2018-05-27 | 0 | 18.8 | 10.6 |
| 2018-05-28 | 0 | 19.4 | 4.5 |
| 2018-05-29 | 5 | 18 | 10.3 |
| 2018-05-30 | 0 | 14.5 | 4.5 |
| 2018-05-31 | 0 | 14.7 | 3.8 |
| 2018-06-01 | 0.9 | 12.8 | 0.2 |
| 2018-06-02 | 0 | 12.2 | 1.2 |
| 2018-06-03 | 0 | 12.3 | -2.5 |
| 2018-06-04 | 0 | 12.5 | 2.1 |
| 2018-06-05 | 1 | 9 | 3 |
| 2018-06-06 | 0 | 12 | 0.5 |
| 2018-06-07 | 1.4 | 12 | 5.2 |
| 2018-06-08 | 1 | 12 | 3.1 |
| 2018-06-09 | 0 | 12.5 | 4.1 |
| 2018-06-10 | 0 | 14 | 3.9 |
| 2018-06-11 | 0.2 | 15.1 | 6.4 |
| 2018-06-12 | 0.2 | 11.6 | -0.5 |
| 2018-06-13 | 4.6 | 8.4 | 6.4 |
| 2018-06-14 | 1 | 9.1 | 4.6 |
| 2018-06-15 | 0.1 | 9.4 | 0.6 |
| 2018-06-16 | 0 | 10.5 | -2.7 |
| 2018-06-17 | 0 | 14 | 1.9 |
| 2018-06-18 | 0 | 16.6 | 6.3 |
| 2018-06-19 | 0 | 19.8 | 7.4 |
| 2018-06-20 | 0 | 18 | 8.2 |
| 2018-06-21 | 0 | 19.6 | -0.4 |
| 2018-06-22 | 0 | 14 | 1.9 |
| 2018-06-23 | 0 | 16 | 1 |
| 2018-06-24 | 0 | 11.5 | 1.7 |
| 2018-06-25 | 0 | 11.8 | -4.5 |
| 2018-06-26 | 0 | 14 | 3.6 |
| 2018-06-27 | 0 | 14.5 | 5.8 |
| 2018-06-28 | 0.4 | 18.6 | 6 |
| 2018-06-29 | 0 | 14.7 | 6.3 |
| 2018-06-30 | 0 | 18.2 | 6 |
| 2018-07-01 | 0 | 19.7 | 7.6 |
| 2018-07-02 | 0 | 11.1 | 8.7 |
| 2018-07-03 | 0 | 9.2 | 3 |
| 2018-07-04 | 0 | 6.3 | -5.6 |
| 2018-07-05 | 0 | 8.6 | -0.6 |
| 2018-07-06 | 0 | 11.4 | 3.2 |
| 2018-07-07 | 0 | 10 | 0.8 |
| 2018-07-08 | 9 | 9.6 | 5.6 |
| 2018-07-09 | 1 | 10.5 | 4.6 |
| 2018-07-10 | 0 | 10.1 | 6.3 |
| 2018-07-11 | 0 | 13 | 0.6 |
| 2018-07-12 | 0 | 13.4 | 6.4 |
| 2018-07-13 | 0 | 17 | 6.6 |
| 2018-07-14 | 0 | 16.4 | 6 |
| 2018-07-15 | 0 | 19.2 | 9 |
| 2018-07-16 | 0 | 13 | 1 |
| 2018-07-17 | 0 | 12.4 | -2 |
| 2018-07-18 | 0.4 | 7.4 | 5.2 |
| 2018-07-19 | 0 | 9.6 | 5.2 |
| 2018-07-20 | 0 | 9.4 | 8.2 |
| 2018-07-21 | 11.8 | 7.6 | 4.7 |
| 2018-07-22 | 2 | 14.6 | -1.2 |
| 2018-07-23 | 0 | 11.3 | 1 |
| 2018-07-24 | 0 | 10.5 | -3.8 |
| 2018-07-25 | 0 | 11.8 | -0.5 |
| 2018-07-26 | 0 | 13.6 | -2.8 |
| 2018-07-27 | 0 | 15.4 | -0.9 |
| 2018-07-28 | 0 | 16.2 | 5.5 |
| 2018-07-29 | 0 | 11.4 | 6.1 |
| 2018-07-30 | 0 | 9.2 | 3 |
| 2018-07-31 | 0 | 11.9 | 2.7 |
| 2018-08-01 | 0 | 12.5 | -5.1 |
| 2018-08-02 | 0 | 13.6 | 0.5 |
| 2018-08-03 | 0 | 18.4 | -1.8 |
| 2018-08-04 | 0 | 13.5 | 2.4 |
| 2018-08-05 | 0 | 18.5 | 1.1 |
| 2018-08-06 | 0 | 20.2 | 9 |
| 2018-08-07 | 0 | 21.4 | 5.3 |
| 2018-08-08 | 6 | 16 | 10.2 |
| 2018-08-09 | 9 | 11 | 7.1 |
| 2018-08-10 | 0 | 11.2 | 3.5 |
| 2018-08-11 | 0 | 11.5 | -1.6 |
| 2018-08-12 | 0 | 15.7 | -3 |
| 2018-08-13 | 0 | 16.8 | 5 |
| 2018-08-14 | 3 | 12.2 | 4.4 |
| 2018-08-15 | 1 | 13.6 | 5.2 |
| 2018-08-16 | 0 | 16.4 | -0.6 |
| 2018-08-17 | 0 | 20 | 10 |
| 2018-08-18 | 0 | 20.6 | 12.6 |
| 2018-08-19 | 0 | 14.2 | 6.5 |

**Fig. S2.** Daily minimum and maximum air temperatures (°C) and rainfall (mm) at the CONICET Bahía Blanca from February 2018 to February 2019.
