## Supplementary material for "Ecological fitness cost associated with the AHAS Trp574Leu mutation in feral *Raphanus sativus*": Fig. S3 (from another version of this preprint: version 2, 2020-08-31)

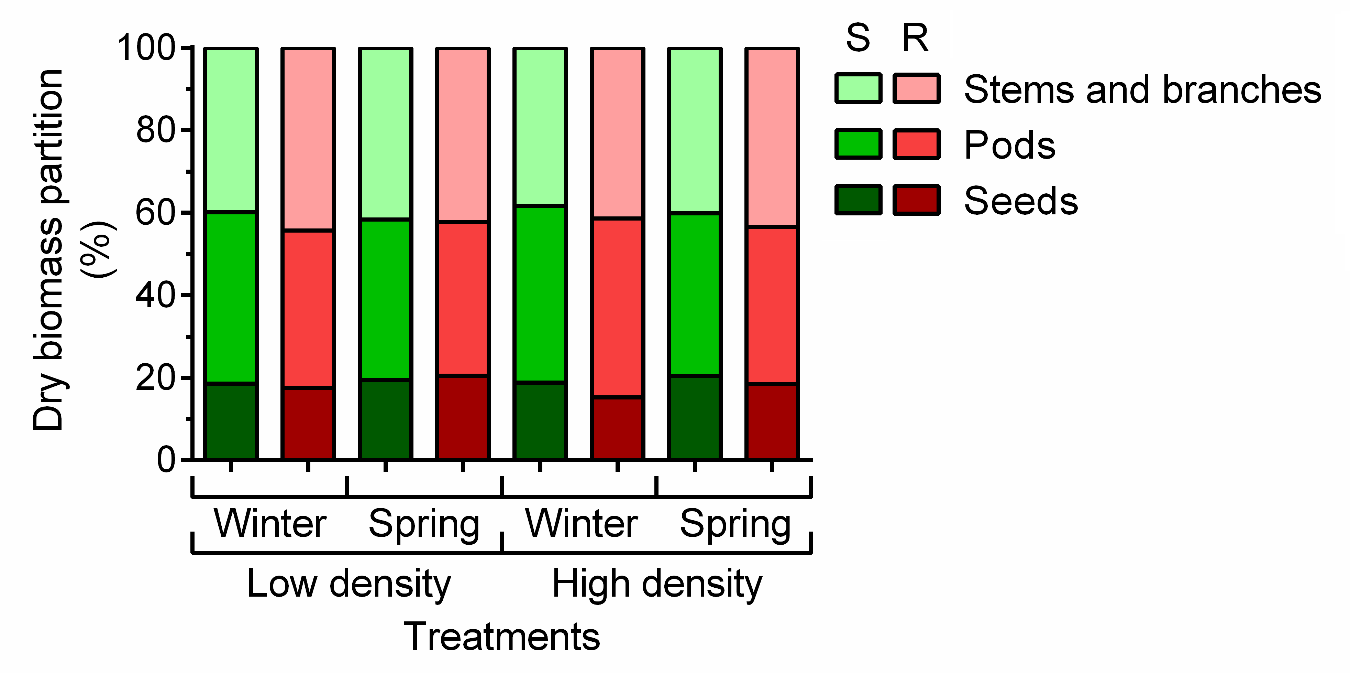

**Figure S3**. Partition of above-ground dry biomass (100 %), in stems and branches, pods and seeds in AHAS herbicide–resistant Trp475Leu (R) and susceptible wild-type (S) feral *Raphanus sativus* grown under wheat competition conditions at two weed densities, low (4 plants m^-2^) and high (12 plants m^-2^), in two planting date (winter and spring). Values are mean of 16 plants (4 blocks x 4 plants block^-1^).
