## Supplementary material for "Ecological fitness cost associated with the AHAS Trp574Leu mutation in feral *Raphanus sativus*": Fig. S4 (from another version of this preprint: version 2, 2020-08-31)

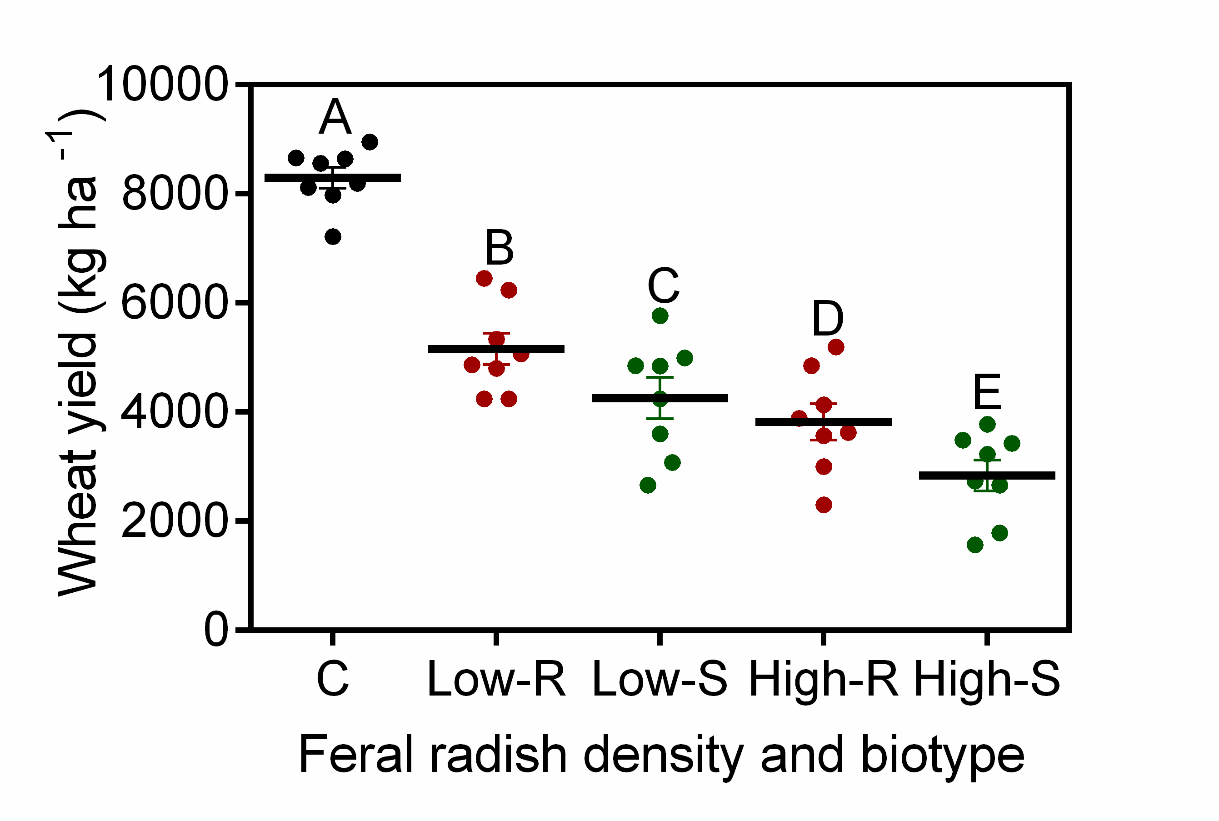

**Figure S4**. Wheat yield under interference of two densities, low (4 plants m^-2^) and high (12 plants m^-2^), of AHAS herbicide-resistant Trp475Leu (R) and susceptible wild-type (S) feral *Raphanus sativus* biotypes. Data are based in two planting date. Horizontal and vertical lines represent mean ± estándar error. Each point indicates the average of three repetitions.
