## Supplementary material for "Ecological fitness cost associated with the AHAS Trp574Leu mutation in feral *Raphanus sativus*": Table S1 (from another version of this preprint: version 2, 2020-08-31)

**Table S1**. A binomial model (two categories: germinated and non-germinated) of effects of light, pericarp treatment (seeds inside intact pods and isolated seeds), temperature, biotype (R and S), storage time (0, 2, 6, and 12 months), and all the interactions on germination of feral *Raphanus sativus* stored dry under laboratory conditions for 12 months. Significant effects (*P* ≤ 0.05) are in bold.

| Effects | NUM_DF_ | DEN_DF_ | *F* | *P* |
| --- | --- | --- | --- | --- |
| Light (L) | 1 | 288 | **102.68** | **<.0001** |
| Pericarp treatment (P) | 1 | 288 | **77.57** | **<.0001** |
| Temperature (Te) | 2 | 288 | **31.05** | **<.0001** |
| Biotype (B) | 1 | 288 | **6.70** | **0.0101** |
| Time (Ti) | 3 | 288 | 1.93 | 0.1254 |
| L x P | 1 | 288 | **23.97** | **<.0001** |
| L x Te | 2 | 288 | 1.74 | 0.1775 |
| L x B | 1 | 288 | 1.96 | 0.1629 |
| L x Ti | 3 | 288 | 0.11 | 0.9550 |
| P x Te | 2 | 288 | 1.19 | 0.3072 |
| P x B | 1 | 288 | 0.02 | 0.8842 |
| P x Ti | 3 | 288 | 1.12 | 0.3423 |
| Te x B | 2 | 288 | 0.06 | 0.9412 |
| Te x Ti | 6 | 288 | 1.31 | 0.2542 |
| B x Ti | 3 | 288 | 0.10 | 0.9611 |
| L x P x Te | 2 | 288 | 0.80 | 0.4515 |
| L x P x B | 1 | 288 | 0.00 | 0.9502 |
| L x P x Ti | 3 | 288 | 0.30 | 0.8249 |
| L x Te x B | 2 | 288 | 0.26 | 0.7675 |
| L x Te x Ti | 6 | 288 | 1.14 | 0.3364 |
| L x B x Ti | 3 | 288 | 0.35 | 0.7884 |
| P x Te x B | 2 | 288 | 0.26 | 0.7733 |
| P x Te x Ti | 6 | 288 | 0.18 | 0.9833 |
| P x B x Ti | 3 | 288 | 0.56 | 0.6444 |
| Te x B x Ti | 6 | 288 | 0.14 | 0.9901 |
| L x P x Te x B | 2 | 288 | 1.36 | 0.2580 |
| L x P x Te x Ti | 6 | 288 | 0.35 | 0.9076 |
| L x P x B x Ti | 3 | 288 | 0.09 | 0.9644 |
| L x Te x B x Ti | 6 | 288 | 0.17 | 0.9839 |
| P x Te x B x Ti | 6 | 288 | 0.47 | 0.8270 |
| L x P x Te x B x Ti | 6 | 288 | 0.89 | 0.5003 |
