## Supplementary material for "Ecological fitness cost associated with the AHAS Trp574Leu mutation in feral *Raphanus sativus*": Table S2 (from another version of this preprint: version 2, 2020-08-31)

**Table 2S**. *ANOVA* results for effects of water condition (watered and non-watered soil), biotype (R and S), pericarp treatment (isolated seeds and seeds inside pod) and their interactions on seedling emergence of feral *Raphanus sativus* incubated under outdoor conditions. Effects were evaluated at five time from the beginning to the end of the experiment (1, 3, 6, 9 and 12 months). Significant effects (*P* ≤ 0.05) are in bold.

|  |  | 1-month | | 3-month | | 6-month | | 9-month | | 12-month | |
| --- | --- | --- | --- | --- | --- | --- | --- | --- | --- | --- | --- |
| Effects | DF_NUM_/DF_DEN_ | *F* | *P* | *F* | *P* | *F* | *P* | *F* | *P* | *F* | *P* |
| Water condition (W) | 1/40 | **68.21** | **<.0001** | **14.57** | **0.0005** | **12.26** | **0.0012** | 2.70 | 0.1080 | **6.32** | **0.0160** |
| Biotype (B) | 1/40 | **21.13** | **<.0001** | **21.82** | **<.0001** | **15.74** | **0.0003** | **9.40** | **0.0039** | **7.31** | **0.0100** |
| Pericarp treatment (P) | 1/40 | **424.66** | **<.0001** | **243.09** | **<.0001** | **184.56** | **<.0001** | **84.50** | **<.0001** | **68.21** | **<.0001** |
| W x B | 1/40 | **6.50** | **0.0148** | **13.27** | **0.0008** | **10.63** | **0.0023** | 2.27 | 0.1395 | 2.61 | 0.1138 |
| W x P | 1/40 | **23.34** | **<.0001** | **11.87** | **0.0014** | **9.13** | **0.0044** | **4.33** | **0.0439** | **6.00** | **0.0188** |
| B x P | 1/40 | 0.07 | 0.7979 | 1.54 | 0.2217 | 2.88 | 0.0974 | 1.24 | 0.2720 | 1.55 | 0.2207 |
| W x B x P | 1/40 | 0.68 | 0.4163 | 0.21 | 0.6489 | 0.34 | 0.5649 | 1.03 | 0.3157 | 0.76 | 0.3882 |
