## Supplementary material for "Ecological fitness cost associated with the AHAS Trp574Leu mutation in feral *Raphanus sativus*": Table S3 (from another version of this preprint: version 2, 2020-08-31)

**Table S3**. *ANOVA* results for effects of biotype (R and S), pericarp treatment (isolated seeds and seeds inside pod) and its interaction on seedling emergence of feral *Raphanus sativus* incubated under outdoor conditions in watered and non-watered soils. Effects were evaluated at five time from the beginning to the end of the experiment (1, 3, 6, 9 and 12 months). Significant effects (*P* ≤ 0.05) are in bold.

|  |  | 1-month | | 3-month | | 6-month | | 9-month | | 12-month | |
| --- | --- | --- | --- | --- | --- | --- | --- | --- | --- | --- | --- |
| Effects | DF_NUM_/DF_DEN_ | *F* | *P* | *F* | *P* | *F* | *P* | *F* | *P* | *F* | *P* |
| Watered |  |  |  |  |  |  |  |  |  |  |  |
| Biotype (B) | 1/20 | 3.89 | 0.06 | 0.94 | 0.34 | 0.50 | 0.49 | 3.50 | 0.08 | 1.25 | 0.28 |
| Pericarp treatment (P) | 1/20 | **234.77** | **<.0001** | **127.82** | **<.0001** | **101.44** | **<.0001** | **73.91** | **<.0001** | **35.86** | **<.0001** |
| B x P | 1/20 | 0.32 | 0.58 | 0.54 | 0.47 | 1.14 | 0.30 | 0.05 | 0.83 | 0.18 | 0.68 |
| Non-watered |  |  |  |  |  |  |  |  |  |  |  |
| Biotype (B) | 1/20 | **15.93** | **0.0008** | **23.16** | **0.0001** | **17.44** | **0.0005** | **6.20** | **0.0217** | **6.19** | **0.0218** |
| Pericarp treatment (P) | 1/20 | **199.51** | **<.0001** | **121.31** | **<.0001** | **92.44** | **<.0001** | **38.12** | **<.0001** | **38.18** | **<.0001** |
| B x P | 1/20 | 0.33 | 0.57 | 0.88 | 0.36 | 1.58 | 0.22 | 1.17 | 0.29 | 1.34 | 0.26 |
