## Supplementary material for "Ecological fitness cost associated with the AHAS Trp574Leu mutation in feral *Raphanus sativus*": Table S4 (from another version of this preprint: version 2, 2020-08-31)

Table S4. *ANOVA* results for effects of planting date (winter and spring), biotype (R and S), density (4 and 12 plants m^-2^), their interactions and block inside planting date on fitness traits of AHAS herbicide-resistant Trp475Leu and susceptible wild-type feral *Raphanus sativus* biotypes grown in wheat competition under field conditions. Significant effects (*P* ≤ 0.05) are in bold.

|  |  | Plant height | | Branch number | | Total above-ground dry biomass plant^-1^ | | Pod number plant^-1^ | | Seed number pod^-1^ | | Seed number plant^-1^ | | Seed weight | | Plant yield | | Biomass allocation to reproductive structures | |
| --- | --- | --- | --- | --- | --- | --- | --- | --- | --- | --- | --- | --- | --- | --- | --- | --- | --- | --- | --- |
| Efectos | DF_N_/DF_D_ | *F* | *P* | *F* | *P* | *F* | *P* | *F* | *P* | *F* | *P* | *F* | *P* | *F* | *P* | *F* | *P* | *F* | *P* |
| Planting date (P) | 1/31 | **45,36** | **<0,0001** | 0,52 | 0,48 | 0,25 | 0,62 | 1,13 | 0.30 | **53,45** | **<0,0001** | **20,05** | **0,0003** | **84,62** | **<0,0001** | 3,16 | 0,09 | **29.08** | **<.0001** |
| Bloque | 6/31 | 2,49 | 0,06 | 0,25 | 0,95 | 2,32 | 0,08 | 1,11 | 0.39 | 0,04 | 0,99 | 1,33 | 0,29 | 1,19 | 0,36 | 2,38 | 0,07 | 2.28 | 0.08 |
| Biotype (B) | 1/31 | 0,24 | 0,63 | 1,85 | 0,19 | **37,00** | **<0,0001** | **16,20** | **0.0008** | 1,07 | 0,31 | **27,50** | **<0,0001** | **19,00** | **0,0004** | **51,05** | **<0,0001** | **11.49** | **0.0033** |
| Density (D) | 1/31 | 0,08 | 0,78 | 0,55 | 0,47 | **14,45** | **0,0013** | **10,57** | **0.0044** | 0,35 | 0,56 | **17,06** | **0,0006** | 1,14 | 0,30 | **17,19** | **0,0006** | 3.94 | 0.06 |
| P x B | 1/31 | 0,25 | 0,62 | 2,60 | 0,12 | 1,08 | 0,31 | 0,44 | 0.51 | 2,18 | 0,16 | 0,68 | 0,42 | 0,90 | 0,36 | 0,65 | 0,43 | **5.28** | **0.0337** |
| P x D | 1/31 | 0,84 | 0,37 | 0,28 | 0,60 | 0,01 | 0,92 | 0,00 | 0.97 | 0,66 | 0,43 | 0,09 | 0,77 | 0,72 | 0,41 | 0,01 | 0,92 | 0.31 | 0.5826 |
| B x D | 1/31 | 0,04 | 0,84 | 1,17 | 0,29 | 0,00 | 0,99 | 1,94 | 0.18 | 1,12 | 0,30 | 1,01 | 0,33 | 0,09 | 0,77 | 0,49 | 0,50 | **12.47** | **0.0024** |
| P x B x D | 1/31 | 0,26 | 0,62 | 0,02 | 0,88 | 0,18 | 0,68 | 0,00 | 0.96 | 0,02 | 0,87 | 0,19 | 0,67 | 1,90 | 0,18 | 0,06 | 0,82 | 0.03 | 0.8719 |
